# Shared mechanisms of auditory and non-auditory vocal learning in the songbird brain

**DOI:** 10.1101/2021.12.09.471883

**Authors:** James N. McGregor, Abigail Grassler, Paul Jaffe, Amanda Jacob, Michael S. Brainard, Samuel J. Sober

## Abstract

Songbirds and humans share the ability to adaptively modify their vocalizations based on sensory feedback. Prior studies have focused primarily on the role that auditory feedback plays in shaping vocal output throughout life. In contrast, it is unclear whether and how non-auditory information drives vocal plasticity. Here, we first used a reinforcement learning paradigm to establish that non-auditory feedback can drive vocal learning in adult songbirds. We then assessed the role of a songbird basal ganglia-thalamocortical pathway critical to auditory vocal learning in this novel form of vocal plasticity. We found that both this circuit and its dopaminergic inputs are necessary for non-auditory vocal learning, demonstrating that this pathway is not specialized exclusively for auditory-driven vocal learning. The ability of this circuit to use both auditory and non-auditory information to guide vocal learning may reflect a general principle for the neural systems that support vocal plasticity across species.

## Introduction

A fundamental goal of neuroscience is to understand how the brain uses sensory feedback to drive adaptive changes in motor output^1,2^. Human speech is a prime example of a sensory-guided behavior, and humans are among the few species that use auditory feedback from their own vocalizations to shape vocal output^3^. This reliance on sensory feedback for speech production is lifelong: loss of hearing impairs both speech development and vocal production in adulthood, and adult speakers rely heavily on auditory signals to calibrate their vocal acoustics^4–7^. Accordingly, studies of the neurobiology of speech have focused on the specialized neural pathways that process auditory feeback^8^. In contrast, it is unclear whether the brain uses non-auditory sensory input to regulate vocal production, although studies demonstrating that humans use non-auditory (somatosensory) signals to calibrate jaw movements suggest that this might be the case^9,10^.

We address how the brain processes different sources of sensory feedback to guide vocal behavior by using a model system ideally suited for the study of vocal learning, the Bengalese finch. Like humans, songbirds rely on auditory signals to precisely calibrate their vocal output throughout life^11–14^. Also similar to humans, songbirds have evolved specialized neural pathways for vocal learning, allowing the precise interrogation of the brain mechanisms of song plasticity^8,15^. However, prior research on this brain network has focused almost exclusively on the role of auditory feedback. These studies have revealed that songbird brains have a basal ganglia-thalamocortical circuit, the Anterior Forebrain Pathway (AFP), that is required for auditory-guided vocal learning but not vocal production (Fig. 1a)^16–19^. For example, lesions of LMAN (the output nucleus of the AFP) prevent adult vocal plasticity in response to perturbations of auditory feedback^16,20,21^. Also, lesions or manipulations of dopaminergic input into Area X (the basal ganglia nucleus of the AFP) impair adult vocal learning in response to the pitch-contingent delivery of aversive auditory stimuli (white noise bursts)^22–24^. Although recent work has demonstrated that the songbird AFP receives anatomical projections from brain regions that process non-auditory sensory information^25^, it remains unknown whether non-auditory information is processed by this circuit to drive vocal learning.

**Figure 1.**
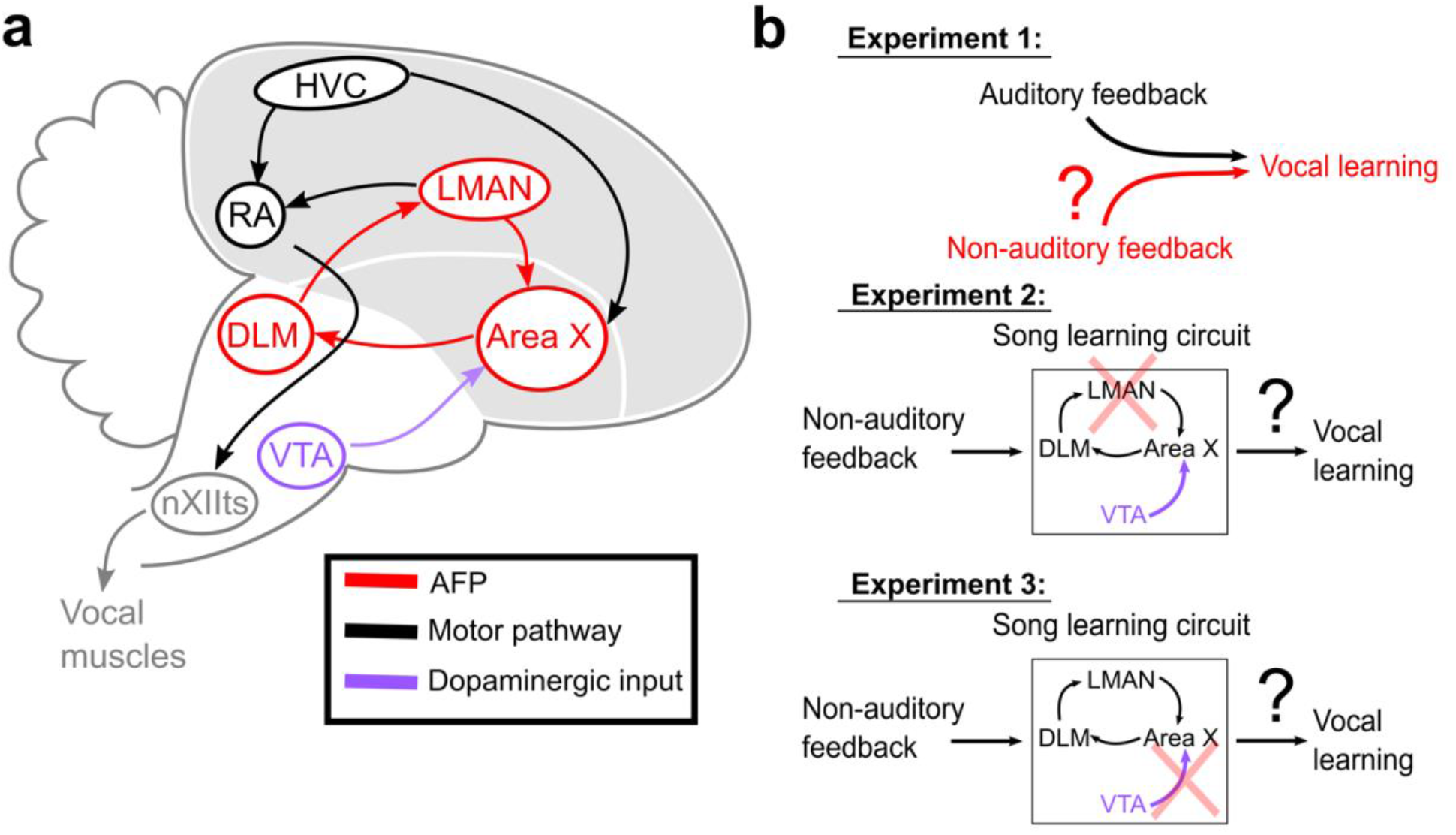
(**a**) Songbird brain circuitry. Brain nuclei of the motor pathway – the neural circuit for vocal production – are black. Brain nuclei of the Anterior Forebrain Pathway (AFP) – the neural circuit for vocal learning – are red. VTA (purple) provides dopaminergic input into Area X, the basal ganglia nucleus of the AFP. (**b**) The three primary hypotheses tested in this paper. In the first set of experiments, we tested whether non-auditory input can drive adaptive changes to adult song (Experiment 1). In the second set of experiments, we assessed the necessity of LMAN for non-auditory vocal learning (Experiment 2). In the third set of experiments, we tested the necessity of dopaminergic projections to Area X for non-auditory vocal learning (Experiment 3).

We performed a series of three experiments (Fig. 1b) to investigate whether and how the brain uses non-auditory sensory feedback to guide vocal learning. We first tested whether adult songbirds can adaptively modify specific elements of their song structure in response to non-auditory feedback (Fig. 1b, Experiment 1). We used non-auditory stimuli (mild cutaneous electrical stimulation), which we delivered during ongoing song performance, to differentially reinforce the acoustics (fundamental frequency, or “pitch”) of specific song elements, or “syllables”. In separate experiments, we tested birds using auditory stimuli consisting brief playbacks of white noise, a well-established paradigm for driving changes in pitch in adult songbirds^22,26,27^. Delivering non-auditory and auditory stimuli on the same schedule therefore allowed us to directly compare how different sensory modalities affect vocal behavior. We next assessed the neural circuit mechanisms underlying non-auditory vocal learning by determining the necessity of LMAN (the output nucleus of the AFP) for non-auditory learning (Fig. 1b, Experiment 2). Finally, we assessed the role of dopaminergic neural circuitry in non-auditory vocal learning by performing selective lesions of dopaminergic input to Area X (Fig. 1b, Experiment 3).

## Results

### Non-auditory feedback can drive adult songbird vocal learning

We tested whether non-auditory feedback can drive vocal learning (Fig. 1b, Experiment 1) by providing mild, pitch-contingent cutaneous stimulation through a set of wire electrodes on the scalps of adult songbirds. Before initiating cutaneous stimulation training, we continuously recorded song without providing any feedback for three days (baseline) (Fig. 2a). Every day, songbirds naturally produce many renditions of song, which consist of repeated patterns of unique vocal gestures, called syllables (Fig. 2b, top). For one “target” syllable in each experimental subject, we quantified rendition-to-rendition variability in the fundamental frequency of each occurrence of this syllable on the final baseline day (Fig. 2b, top). To differentially reinforce the pitch of a target syllable, we determined a range of pitches within this baseline distribution (either all pitches above the 20th percentile or all pitches below the 80th percentile), and then triggered the delivery of cutaneous stimulation in real time (within 40 ms of syllable onset) when the pitch of the target syllable fell within this range (Fig. 2b, bottom). We performed this pitch-contingent cutaneous stimulation training continuously for three days. Note that the birds could choose not to sing in order to avoid triggering any cutaneous stimulation, and we carefully monitored animal subjects for any signs of distress (see Methods).

**Figure 2.**
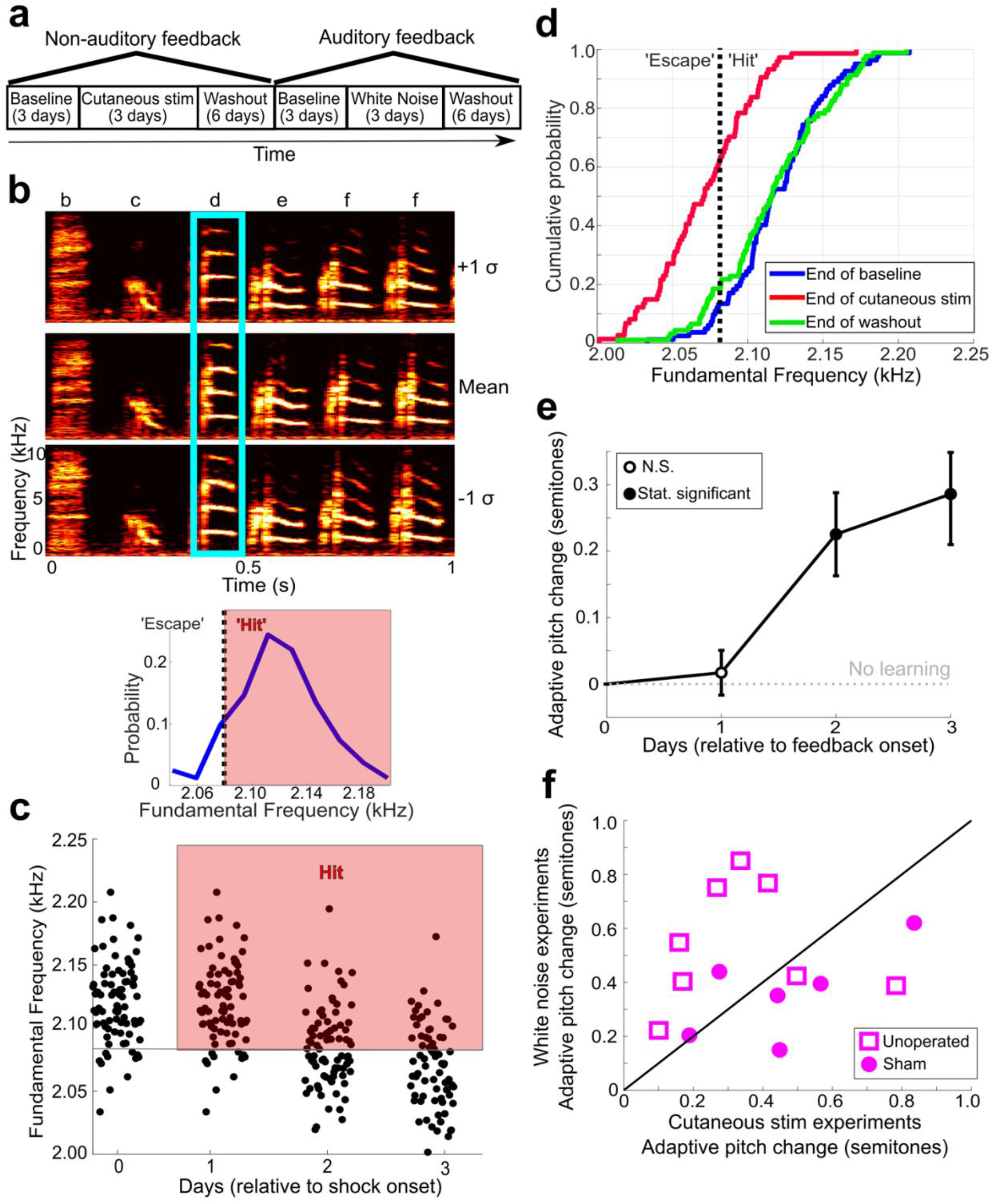
Non-auditory feedback drives vocal learning. (**a**) Timeline of vocal learning experiments in individual birds. The order of the auditory vs non-auditory experiments was randomized across birds. (**b**) (Top) Spectrograms and song syllables (labeled b-f) including target syllable (“d”). (Bottom) baseline pitch distribution and pitch threshold. Cutaneous stimulation was provided during renditions of the target syllable above a chosen pitch threshold (“hit”). (**c**) Each dot represents the pitch of one rendition of the target syllable. Renditions in the “hit” range rapidly triggered a cutaneous stimulation (within 40 ms of syllable onset). (**d**) CDF plot showing the probability a value of pitch from a distribution falls at or below the value on the x-axis. The pitch distribution at the end of cutaneous stimulation training was significantly greater than baseline (2-sample KS test, p=1.178e-12). End of washout distribution was not significantly different from baseline (2-sample KS test, p=0.606). Panels B-D show data from the same experiment. (**e**) Adaptive pitch change (in semitones) of the target syllables during cutaneous stimulation training, grouped across 13 experiments. The mean change during training was significantly greater than baseline (probability of resampled mean pitch on all three training days 2 and 3 lesser than or equal to zero was P_boot_<0.0010, indicated by filled circles). (**f**) Learning magnitudes (adaptive pitch change by end of training) in individual birds that underwent both white noise and cutaneous stimulation training (n=14). No significant difference in learning magnitudes during cutaneous stimulation training vs during white noise training (paired t-test, p=0.313).

For example, in one experiment (shown in Fig. 2a-d), cutaneous stimulation was triggered on every rendition of the target syllable that had a pitch above 2.13 kHz (the 20th percentile of the baseline distribution) for three days. In this example experiment, the bird gradually changed the pitch of the targeted syllable downwards (the adaptive direction), such that cutaneous stimulation was triggered less frequently (Fig. 2c). In other experiments where the adaptive direction of pitch change is upwards, we triggered cutaneous stimulation whenever the target syllable pitch was below the 80th percentile of this distribution. In the example experiment, at the start of the first day of cutaneous stimulation training, 80% of syllable renditions resulted in cutaneous stimulation and 20% of syllable renditions resulted in escapes. On the third (final) day of cutaneous stimulation training, escapes occurred on over 60% of target syllable renditions and the entire distribution of pitches had changed significantly in the adaptive direction, indicating that a significant amount of vocal learning occurred in this example experiment (Fig. 2d; 2-sample KS test to assess the difference between baseline and end of cutaneous stimulation training, p = 1.1776e-12). We then stopped triggering cutaneous stimulation and continued to record unperturbed song for six additional days (washout). After six days of washout, there was no significant difference between the distribution of target syllable pitches at the end of washout compared to baseline (Fig. 2d; 2-sample KS test, p = 0.606). For analysis of washout across all experiments, see Figure 2-Figure Supplement 1.

In order to assess whether non-auditory feedback is sufficient to drive vocal learning across multiple songbirds, we first measured the adaptive pitch change (in semitones) for each individual experiment. Semitones provide a normalized measure of pitch change such that a one semitone change corresponds to a roughly 6% change in the absolute frequency of an acoustic signal (see Equation 1 in Methods). We employed a hierarchical bootstrap approach to measure SEM and assess significance (see Methods)^28,29^ since this method more accurately quantifies the error in hierarchical data (e.g., many renditions of a target syllable collected across multiple birds). We found that the mean pitch (in semitones) of the target syllables showed a significant, adaptive change from baseline on days two and three of cutaneous stimulation training (Fig. 2e; probability of resampled mean pitch on cutaneous stimulation training days 2 and 3 lesser than or equal to zero was P_boot_ < 0.0010, limit due to resampling 10^4^ times). This demonstrates that non-auditory feedback is sufficient to drive vocal learning in adult songbirds. In all individual experiments where an upwards pitch change resulted in less frequent triggering of cutaneous stimulation, the birds changed their pitch in the adaptive (upward) direction, and in all experiments where a downwards pitch change resulted in less frequent triggering of cutaneous stimulation, the birds changed their pitch in the adaptive (downward) direction (Figure 2-Figure Supplement 2a).

To compare vocal learning in response to different sources of sensory feedback (auditory and non-auditory), we performed multiple learning experiments -one cutaneous stimulation and one white noise -in 8 out of the 12 individual birds from this data set (Fig. 2a). We randomized the order of white noise training and cutaneous stimulation training for the birds who underwent both training paradigms. We also included 6 sham operated birds from a later set of experiments in this particular analysis. We did so because the sham operated birds had intact song systems and underwent both cutaneous stimulation and white noise training.

Consistent with prior studies^20,22,27^, by the end of white noise training, the adaptive pitch change (in semitones) across all white noise experiments performed in unoperated birds (birds who had wire electrodes surgically implanted but received no invasive brain procedures like sham operations) was significantly greater than baseline (zero) (Figure 2-Figure Supplement 3a; probability of resampled mean pitch on all three cutaneous stimulation training days lesser than or equal to zero was P_boot_ < 0.0010). In the separate experimental group of birds that underwent sham operations, we also observed significant adaptive pitch changes in response to white noise bursts, as expected (Figure 2-Figure Supplement 3b, probability of resampled mean pitch on all three cutaneous stimulation training days lesser than or equal to zero was P_boot_ < 0.0010). There was individual variability in learning magnitudes (adaptive pitch change at the end of training) during cutaneous stimulation and white noise experiments (Fig. 2f). However, we found no systematic differences between learning magnitude during cutaneous stimulation training and the learning magnitude during white noise training (Fig. 2f; paired t-test, p = 0.313). These results suggest that non-auditory stimuli can drive vocal learning as effectively as auditory stimuli.

To confirm that cutaneous stimulation was truly non-auditory and did not produce any acute changes to vocal output, we measured the pitch of interleaved “catch” trials, where cutaneous stimulation was randomly withheld (see Methods), on each day of cutaneous stimulation training. For each experiment described in this paper, we normalized the pitch of each catch trial from each day of training to the mean pitch of all trials where cutaneous stimulation was provided. We excluded any experiments where the total number of catch trials was less than 10. In every case, the normalized catch trials did not differ significantly from 1, indicating that the pitch of catch trials were highly similar to trials where cutaneous stimulation was provided (Figure 2-Figure Supplement 4a; t-test, 0.071 < p < 0.997 for each experiment). For comparison, we also performed the same analysis on randomly selected trials from a day of baseline recording, where cutaneous stimulation was not provided on any trials (Figure 2-Figure Supplement 4b). There was no significant difference between this data set and the normalized catch trials (paired t-test, p = 0.339).

### LMAN is required for non-auditory vocal learning

We next investigated the neural circuitry that processes non-auditory feedback to drive vocal learning. To assess whether the AFP is required for non-auditory vocal learning, we measured the effect of lesions of LMAN, the output nucleus of the AFP, on learning magnitude in response to non-auditory feedback (Fig. 1b, Experiment 2). We performed cutaneous stimulation training experiments in the same individual birds before and after bilateral, electrolytic LMAN lesions or sham operations (Fig. 3a, n = 5 birds). To perform cutaneous stimulation training in this group of experiments, we used the same protocol described previously, except we extended the period of cutaneous stimulation training by 1-5 days. During this extended training period, we set a new pitch threshold each morning to drive even greater amounts of learning (“staircase” training, see Methods). In adult songbirds with intact song systems (prelesion), such staircase training drove significant amounts of learning (Fig. 3c).

**Figure 3.**
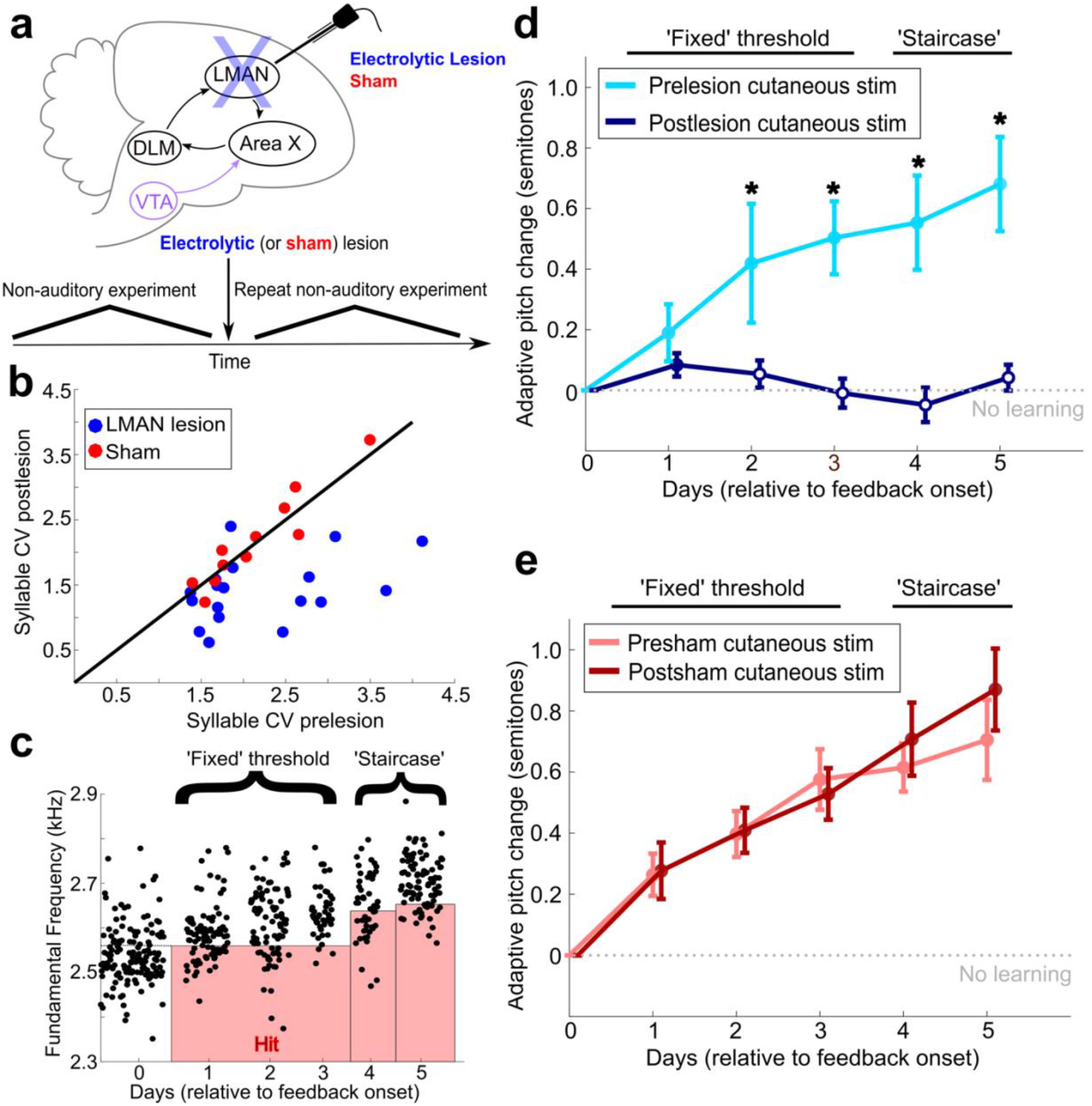
LMAN is required for non-auditory vocal learning. (**a**) Timeline for electrolytic lesions of LMAN and sham operations. (**b**) CV of syllable pitch pre-vs postlesion and pre-vs postsham. LMAN lesions induced a significant reduction in pitch CV, sham operations did not (paired t-tests, p=0.002, p=0.911, respectively) (**c**) Prelesion experiment. Training consisted of three days using a fixed pitch threshold, then additional days where the threshold was changed each morning (“staircase”). Each dot represents the pitch of a rendition of the target syllable. (**d**) Adaptive pitch change (in semitones) during cutaneous stimulation training (n=6 LMAN lesioned birds). Prelesion learning magnitude was significantly greater than baseline (probability of resampled mean pitch on each day of training lesser than or equal to zero was P_boot_<0.0010, indicated by filled circles). Postlesion learning magnitude did not significantly differ from baseline (0.297<P_boot_<0.660 on each of the final four days of training). Prelesion learning magnitude was significantly greater than postlesion learning magnitude (probability of resampled mean pitch of prelesion data on the final 4 days of training lesser than or equal to resampled mean pitch of postlesion data was P_boot_<0.0070, indicated by asterisks). (**e**) Adaptive pitch change during cutaneous stimulation training (n=5 sham operated birds). Learning magnitudes were significantly greater than baseline both pre-and postsham (probability of resampled mean pitch on each day of training lesser than or equal to zero was P_boot_<0.0010, indicated by filled circles). Learning magnitudes pre-vs postsham did not significantly differ (0.120<P_boot_<0.524 on all days of training).

We then lesioned LMAN and performed postlesion white noise training across conditions (LMAN lesion and sham) (Figure 2-Figure Supplement 3b). The efficacy of LMAN lesions was confirmed both by the presence of a characteristic reduction in the trial-to-trial variability of syllable pitch (Fig. 3b and Figure 3-Figure Supplement 1a, LMAN lesions p = 0.002, sham lesions p = 0.911, paired t-tests)^30–32^ and by post-hoc histological measurements (see Methods and Figure 3-Figure Supplement 2). Following LMAN lesions, songbirds did not significantly change the pitch of the target syllable from baseline (zero) (probability of resampled mean pitch on the final four days of cutaneous stimulation training lesser than or equal to zero was P_boot_ > 0.223). In contrast, following sham lesions, birds significantly changed the pitch of the target syllable in the adaptive direction (probability of resampled mean pitch on the final four days of cutaneous stimulation training days lesser than or equal to zero was P_boot_ < 0.0010). This indicates that LMAN lesions induced significant deficits in auditory vocal learning, consistent with previous work that demonstrated that electrolytic LMAN lesions inhibit auditory vocal learning^25^.

LMAN lesions also significantly impaired non-auditory vocal learning. Prelesion, songbirds adaptively changed the pitch of the target syllable away from baseline in response to non-auditory feedback (probability of resampled mean pitch on each day of cutaneous stimulation training lesser than or equal to zero was P_boot_ < 0.0010) (Fig. 3d). Postlesion, non-auditory vocal learning was abolished in those same birds (probability of resampled mean pitch on each of the final four days of training lesser than or equal to zero was 0.297 < P_boot_ < 0.660, where 0.025 < P_boot_ < 0.975 indicates no significant difference, n = 5 birds) (Fig. 3d). Learning magnitude prelesion was significantly greater compared to learning magnitude postlesion (P_boot_ < 0.007 on each of the final four days of training). We observed significant amounts of learning during cutaneous stimulation training in both pre-and post-sham-lesioned datasets (Fig. 3e, for both presham and postsham datasets, the probability of resampled mean pitch on each day of cutaneous stimulation training lesser than or equal to zero was P_boot_ < 0.0010, n = 6 birds). Also, the learning magnitudes during cutaneous stimulation training did not significantly differ in pre-vs postsham datasets (probability of resampled mean pitch of presham data on each day of training lesser than or equal to resampled mean pitch of postlesion data was 0.120 < P_boot_ < 0.524). The amount of pitch change during cutaneous stimulation training for each individual experiment is shown in Supplemental Fig. 2b, c.

We also directly compared the lesion-induced change in learning magnitudes between conditions (LMAN lesion vs sham) (Figure 3-Figure Supplement 1b, c). First, we calculated learning magnitude at the end of the fixed threshold training period across conditions. The lesion-induced change in learning magnitude (post – pre) for LMAN lesioned birds was significantly greater than for sham operated birds (Figure 3-Figure Supplement 1b; 2 sample KS test, p = 0.036). Next, we calculated learning magnitude at the end of the extended “staircase” portion of cutaneous stimulation training across conditions. The lesion-induced change in learning magnitude (post – pre) for LMAN lesioned birds calculated at this time point was also significantly greater than for sham lesioned birds (Figure 3-Figure Supplement 1c; 2 sample KS test, p = 0.004). These results indicate that LMAN is required for non-auditory vocal learning in adult songbirds, indicating that both auditory and non-auditory sensory feedback engage the AFP to drive adaptive changes to song.

## Dopaminergic input to Area X is required for non-auditory vocal learning

We next assessed dopaminergic contributions to non-auditory vocal learning (Fig. 1b, Experiment 3). Learning magnitude during cutaneous stimulation training was assessed before and after bilaterally lesioning dopaminergic projections in Area X, the basal ganglia nucleus of the AFP, in individual songbirds (Fig. 4a, n = 5 birds). Selective lesions of dopaminergic projections in Area X were performed via bilateral 6-OHDA injections in Area X (see Methods), and the effectiveness of the 6-OHDA injections at lesioning dopaminergic innervation in Area X was quantified (Figure 4-Figure Supplement 1). This approach has previously been shown to selectively lesion dopaminergic inputs to Area X without damaging non-dopaminergic cells^22,29^.

**Figure 4.**
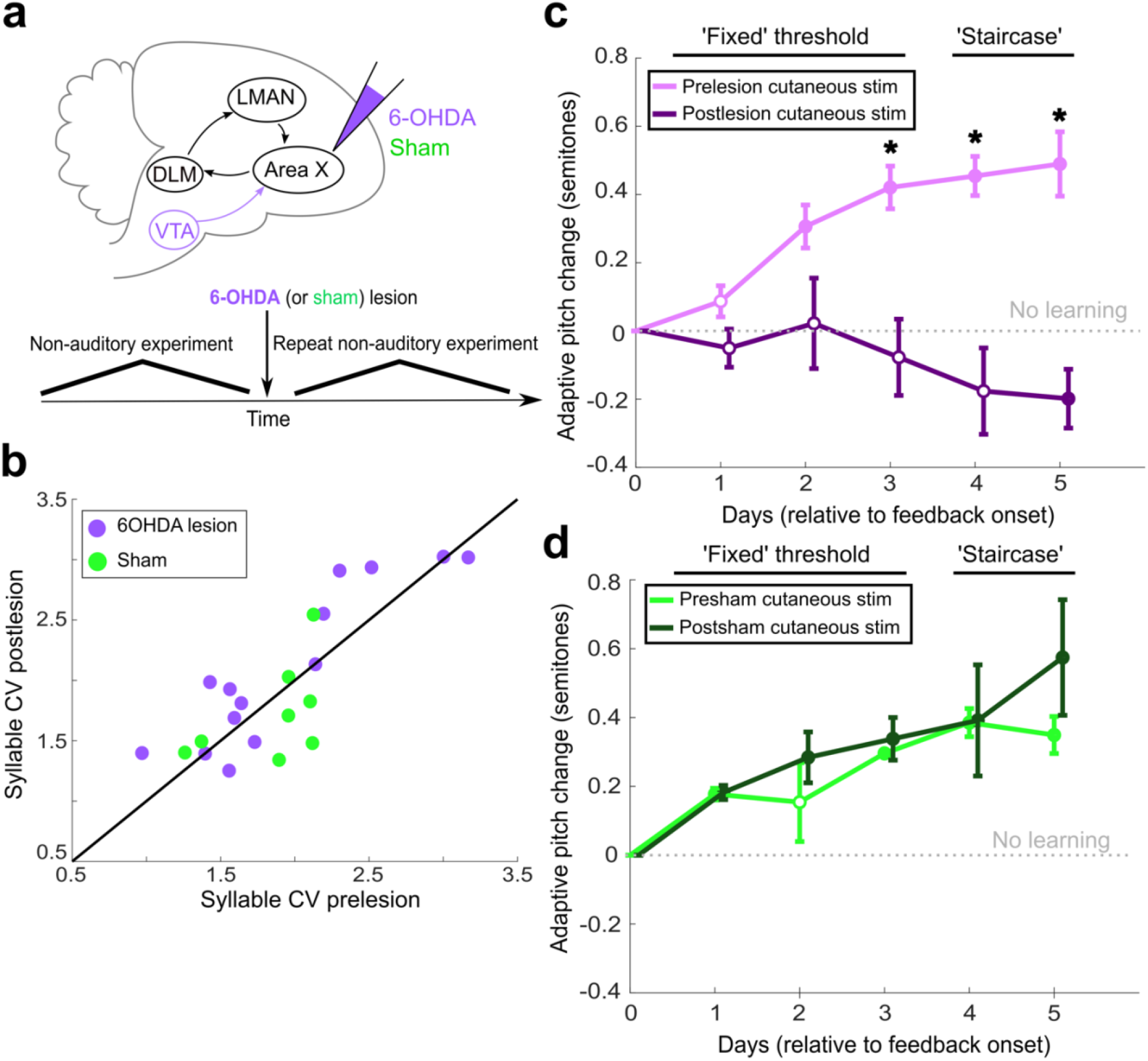
Dopaminergic input to Area X is required for non-auditory vocal learning. (**a**) Timeline for 6-OHDA and saline (sham) injections into Area X. (**b**) CV of syllable pitch pre-vs postlesion and pre-vs postsham. Neither dopamine lesions nor shams induced significant changes in pitch CV (paired t-tests, p=0.397 and p=0.531, respectively). (**c**) Adaptive pitch change (in semitones) during cutaneous stimulation training (n=5 lesioned birds). Prelesion learning magnitude was significantly greater than baseline (probability of resampled mean pitch on each of the final 4 days of training lesser than or equal to zero was P_boot_<0.010, indicated by filled circles). Postlesion learning magnitude did not significantly differ from baseline except for on the final day, when the mean changed in the anti-adaptive direction (P_boot_>0.067 on training days 1-4, P_boot_<0.0010 on training day 5). Prelesion learning magnitude was significantly greater than postlesion learning magnitude (probability of resampled mean pitch from prelesion dataset on each of the final 3 days of training lesser than or equal to resampled mean pitch from postlesion dataset was P_boot_<0.0010, indicated by asterisks). (**d**) Adaptive pitch change (in semitones) during cutaneous stimulation training (n=3 sham birds). Learning magnitudes were significantly greater than baseline both pre-and postsham (probability of resampled mean pitch from presham and postsham datasets on each day other than day 2 of training lesser than or equal to zero was P_boot_<0.0010, indicated by filled circles). Learning magnitudes pre-vs postsham did not significantly differ (0.653<P_boot_<0.931 on all days of training).

We again measured the variability of syllable pitch pre-and postlesion by calculating syllable CV. Dopaminergic lesions in Area X did not induce a significant change in syllable CV (Fig. 4b; paired t-test, p = 0.397). Sham operations also did not induce a significant change in syllable CV (Fig. 4b; paired t-test, p = 0.531). The lesion-induced changes in syllable CV (post -pre) were not significantly different for 6-OHDA lesioned birds than for sham lesioned birds (Figure 3-Figure Supplement 1d; 2 sample KS test, p = 0.054). This finding is consistent with prior work using similar 6-OHDA injections to lesion dopaminergic input to Area X^22^. Prior work has suggested a link between dopamine in songbird AFP and the generation of variability in syllable pitch in adult songbirds^33–35^. It is likely that the dopamine lesion methodology we used, which spares about 50% of the dopaminergic input to Area X^22^, is insufficient to impair dopamine-mediated generation of syllable variability. The result that these dopamine lesions do not alter vocal variability suggests that any learning deficits observed following lesions of AFP circuits are not simply due to decreased pitch variability.

Depletion of dopaminergic input to Area X significantly impaired non-auditory vocal learning. Prelesion, songbirds adaptively changed the pitch of the target syllable during cutaneous stimulation training (probability of resampled mean pitch on each of the final four days of cutaneous stimulation training lesser than or equal to zero was P_boot_ < 0.010) (Fig. 4c). Postlesion, these same birds were not able to adaptively change the pitch of the target syllable during cutaneous stimulation training (probability of resampled mean pitch on each of the first four days of training lesser than or equal to zero was 0.067 < P_boot_ < 0.553. Probability of resampled mean pitch on the final day of training greater than or equal to zero was P_boot_ < 0.0010, n = 5 birds). Learning magnitude prelesion was significantly greater compared to learning magnitude postlesion (probability of resampled mean pitch from prelesion dataset on each of the final 3 days of cutaneous stimulation training lesser than or equal to resampled mean pitch from postlesion dataset was P_boot_ < 0.0010). Both pre-and postsham, songbirds displayed significant amounts of learning during cutaneous stimulation training (Fig. 4d, probability of resampled mean pitch from the presham dataset on each day other than day 2 of cutaneous stimulation training lesser than or equal to zero was P_boot_ < 0.0010. Probability of resampled mean pitch from the postsham dataset on each day of cutaneous stimulation training lesser than or equal to zero was P_boot_ < 0.0010, n = 3 birds). Also, the learning magnitudes during cutaneous stimulation training did not significantly differ pre-vs postsham (probability of resampled mean pitch of presham data on each day of training lesser than or equal to resampled mean pitch of postlesion data was 0.653 < P_boot_ < 0.931). These results demonstrate that dopaminergic input to Area X is required for non-auditory vocal learning. The amount of pitch change during cutaneous stimulation training for each individual experiment is shown in Figure 2-Figure Supplement 2d, e.

## Discussion

Our results demonstrate that non-auditory feedback can drive vocal learning in adult songbirds and that the AFP and its dopaminergic inputs are required for non-auditory vocal learning. We first demonstrated that adult songbirds learn to adaptively change the pitch of their song syllables in response to cutaneous stimulation (Fig. 1b, Experiment 1). We next demonstrated that LMAN, the output nucleus of the AFP, is necessary for the expression of this non-auditory vocal learning (Fig. 1b, Experiment 2). Finally, we showed that dopaminergic input to Area X, the basal ganglia nucleus of the AFP, is necessary for non-auditory vocal learning (Fig. 1b, Experiment 3). These results show that adult vocal learning is not solely dependent on auditory feedback, and that the songbird AFP is not specialized just for processing auditory feedback for vocal learning, as has previously been hypothesized^36^. Instead, these results indicate that the AFP processes auditory feedback as well as non-auditory feedback to drive vocal learning. Prior work has shown that songbird vocal muscles use somatosensory feedback to compensate for experimentally-induced changes in respiratory pressure during song performance^37^. The result that the AFP underlies vocal learning driven by somatosensory signals (cutaneous stimulation) suggests that it could play a role in processing somatosensory information from vocal muscles to guide song performance. Also, the fact that mild cutaneous stimulation is different than the direct proprioceptive feedback from vocal muscles or vocal effectors, yet the AFP still underlies vocal learning in response to cutaneous stimulation, suggests that the AFP can integrate sensory information from a wide variety of sources of sensory feedback, even those not directly produced by vocalizations.

Our findings suggest the importance of neural pathways that convey non-auditory sensory signals to the song system. The neuroanatomical pathways for auditory feedback to enter the AFP are well-characterized. For example, recent work has demonstrated that songbird ventral pallidum (VP) receives input from auditory cortical areas, encodes auditory feedback information, and projects to VTA^38^. This represents a likely pathway by which sensory information from white noise bursts could influence neural activity in VTA, which could then drive changes in the AFP that promote song learning. Comparatively less is known about pathways in the songbird brain that might carry sensory information from cutaneous stimulation to the AFP. The results showing that dopaminergic input to Area X (which originates in the VTA) is necessary for non-auditory vocal learning suggests that pathways for non-auditory information ultimately project to the VTA, where this information could be encoded and transmitted to the AFP to drive learning.

Prior studies have hinted that non-auditory feedback may play an important role in shaping vocalizations in ethological contexts, particularly during development. For example, juvenile songbirds that receive both auditory and visual feedback from live tutors display more accurate copying of tutor songs relative to juvenile songbirds who only receive auditory feedback from their tutors^39^. Also, visual displays from adult song tutors positively reinforce the acquisition of specific song elements in juvenile songbirds^40^, further suggesting an important role for visual signals in social interactions during song learning. Our results that cutaneous stimulation can drive adaptive vocal changes in adult songbirds demonstrates that non-auditory signals, even in the absence of any social cues or other reinforcing sensory signals, can drive vocal learning just as strongly as auditory feedback. Further, our work suggests that the AFP might play a role in processing non-auditory sensory information important to other social behaviors that involve vocal communication, such as courtship, territorial displays, and pair bonding.

It has been hypothesized that a key function of the songbird AFP circuitry is to encode auditory performance error: the evaluation of the match between the auditory feedback the songbirds receive and their internal goal for what their song should sound like (based on their stored memory of the tutor song template)^11,29,41,42^. Some have speculated that white noise bursts are interpreted by the bird as an auditory performance error: an adult songbird expects to hear the auditory feedback from a well-performed song syllable, but instead hears a loud burst of white noise, which it interprets as a poorly-performed song syllable^38,42^. Some evidence supports this hypothesis. For example, pitch-contingent white noise bursts provided during song performance drive adaptive vocal changes^22,27^, but when white noise bursts are provided in non-vocal contexts, such as when a songbird stands on a particular perch (not during song performance), they can positively reinforce place preference^36^. This suggests that white noise is not a generally aversive reinforcement stimulus. In contrast, other reports have suggested that white noise bursts are aversive since white noise bursts are loud and jarring, sound very different than birdsong, and songbirds will adaptively change their vocalizations to avoid triggering white noise bursts as frequently^22,27,43,44^. Although the results of the experiments described here do not prove whether white noise bursts drive learning because the white noise is registered by the birds as a performance error or because the white noise is generally aversive, cutaneous stimulation is an explicit, external sensory stimulus that can drive vocal learning. That the AFP underlies non-auditory learning suggests that the AFP does not solely encode auditory performance error. Instead, the AFP may encode more general information about whether vocal performance resulted in a “good” or “bad” outcome, and it may use this information to drive changes to future motor output.

The numerous analogies between the specialized vocal learning neural circuits that have evolved in songbirds and in humans suggest that our findings may be relevant to understanding the neural circuit mechanisms underlying human speech^3,8,15,45^. Human speech depends on both auditory and non-auditory sensory information to guide learning, yet very little is known about the neural mechanisms for non-auditory vocal learning^46–48^. Our findings show that specialized vocal learning circuitry in songbirds processes non-auditory information to drive vocal learning. We suggest that the analogous vocal circuitry in humans may also underlie non-auditory vocal learning. This neural circuitry in humans may underlie the processing of multimodal sensory signals during social interactions that modulate speech learning^46–48^, or the non-auditory, somatosensory feedback from vocal effectors during speech production^10^.

## Materials and Methods

All subjects were adult (>100 days old) male Bengalese finches (Lonchura striata var. domestica). All procedures were approved by Emory University’s Institutional Animal Care and Use Committee. All singing was undirected (in the absence of a female bird) throughout all experiments.

### Delivery of non-auditory sensory feedback

To deliver non-auditory feedback signals to freely-behaving songbirds during ongoing song performance, we first performed a surgery prior to any experimentation. Stainless steel wires were uninsulated at the tip (2-4 mm) and implanted subcutaneously on the bird’s scalp. In 7 out of all 28 birds used across all experiments performed, wires were implanted intramuscularly in the birds’ necks instead of on their scalps. The wires were soldered onto a custom-made circuit board that, during surgery, was placed on the bird’s skull using dental cement. The circuit was connected to an electric stimulator (A-M Systems Isolated Pulse Stimulator), which produced pitch-contingent electrical currents through the wires implanted on the bird. We set the duration of cutaneous stimulation to 50 ms, which was a long enough duration to overlap with a large portion of the targeted syllable, yet a short enough duration to avoid interfering with following song syllables. We typically set the magnitude of electric current used for producing the shocks to 100-350 μA, which is behaviorally salient (the first few instances of cutaneous stimulation interrupt song), yet subtle enough as to not produce any body movements or signs of distress. Stimulations typically occurred within 20-30 ms of target syllable onset. Acute effects of electrical shock on song structure, such as pitch, amplitude, entropy, or syllable sequence, were assessed to ensure these non-auditory stimuli produced no immediate, systematic, acoustic effects. This ensures that any observed gradual changes to song structure in response to cutaneous stimulation are due to non-auditory learning.

### Vocal learning paradigm and song analysis

Experimental testing of vocal learning was performed by driving adaptive changes in the fundamental frequency (pitch) of song syllables. To do so, we delivered pitch-contingent, non-auditory feedback (mild cutaneous electrical stimulation) to freely-behaving songbirds in real time during song performance. We followed the same experimental protocols as experiments using white noise feedback to drive vocal learning^22,27^, except we used cutaneous stimulation instead of white noise bursts. After surgically implanting the fine-wire electrodes, we recorded song continuously for three days without providing any experimental feedback (cutaneous stimulation or white noise bursts). We refer to this period as “baseline” (Fig. 2a).

On the last (third) day of baseline, we measured the pitch of every rendition of the target syllable sung between 10 a.m. and 12 p.m. We set a fixed pitch threshold based on the distribution of these pitches, such that we would provide sensory feedback only when the pitch of a rendition of the target syllable was above the 20th percentile of the baseline distribution (“hit”), and all renditions outside of this range did not trigger any feedback (“escape”). In this case, an adaptive vocal change would therefore be to change the pitch of the target syllable down, thereby decreasing the frequency of triggering cutaneous stimulation. In other experiments, we triggered feedback on all renditions below the 80th percentile of the baseline pitch distribution. In this case, an adaptive vocal change would be to change the pitch of the target syllable up. For each experiment, we randomly selected which of these two contingencies we employed so we could assess bidirectional adaptations in vocal motor output. In a subset of experiments, we used the 90th percentile and 10th percentile pitch values to set the pitch threshold. Importantly, we also randomly withheld triggering feedback on 10% of syllable renditions, regardless of syllable pitch or the experimental pitch-contingency. This allows us to compare syllable renditions that did or did not result in cutaneous stimulation to assess any acute effects of this form of feedback on syllable structure.

At 10 a.m. on the fourth day of continuous song recording, we began providing pitch-contingent cutaneous stimulation in real time, targeted to specific song syllables sung within a specified range of pitches. We refer to this time period as “cutaneous stimulation training” (Fig. 2 a). We used custom LabVIEW software to continuously record song, monitor song for specific elements indicative of the performance of the target syllable, perform online, rapid pitch calculation, and trigger feedback in real time. The computers running this software were connected to an electric stimulator. When the electric stimulator received input from the LabVIEW software, it would then trigger a 50 ms burst of electric current through the implanted wire electrodes. During cutaneous stimulation training, we continuously recorded song and provided pitch-contingent cutaneous stimulation at the set fixed pitch threshold for three days. During these three days, every time the bird sang within the “hit” range, a mild cutaneous stimulation was immediately triggered.

After three days of cutaneous stimulation training, we stopped providing cutaneous stimulation but continued recording unperturbed song for six additional days. We refer to this period as “washout” (Fig. 2a). During washout, we consistently observed spontaneous pitch restoration back to baseline across all experiments, which is in congruence with results from numerous white noise learning experiments^22,26,27^. This allows for multiple experiments to be performed from similar baseline conditions in the same individual songbird.

In 14 out of all 28 birds used throughout this study, we performed both white noise training and cutaneous stimulation training in the same individual birds (Fig. 2a). After the end of cutaneous stimulation training and six days of washout (when the pitch of the target syllable had restored to baseline levels), we performed the exact same experimental protocol, but we used white noise feedback instead of cutaneous stimulation. We could then compare learning in response to two different sources of sensory feedback in the same individual subject. We also sometimes reversed the order of experimentation by performing white noise experiments first and cutaneous stimulation experiments second. The order of experimentation was randomly decided for each songbird before beginning any white noise or cutaneous stimulation training.

For all LMAN lesion (Fig. 3a) and 6-OHDA lesion experiments (Fig. 4a), we performed a cutaneous stimulation training experiment prelesion. After six days of washout, we then performed surgery to lesion the neural circuit of interest. We then performed another cutaneous stimulation experiment in the same individual bird using the exact same protocol we used prelesion. For all of these lesion cutaneous stimulation experiments, we used the aforementioned cutaneous stimulation training paradigm, but with one slight alteration: we extended the number of days of cutaneous stimulation training and introduced a new methodology for setting the pitch threshold on these extended days of training. We still set a fixed pitch threshold based on analysis of the pitch distribution from the final day of baseline and performed three days of cutaneous stimulation training using this fixed pitch threshold. We refer to this portion of the lesion experiments as “fixed” because the pitch threshold for determining whether a cutaneous stimulation was provided remained the same for all 3 days. Rather than stopping cutaneous stimulation training at this point, we instead continued providing pitch-contingent cutaneous stimulation for an additional 1-5 days. In the morning (at 10 a.m.) on each of these extended days of cutaneous stimulation training, we changed the pitch threshold to the 20th or 80th percentile (consistent with the initial contingency) of the pitch distribution of all renditions of the target syllable sung between 8 A.M. to 9:30 A.M. on that same day. As the bird changed the pitch of the target syllable in the adaptive direction, the new pitch thresholds continued to be set further and further in the adaptive direction to drive greater amounts of learning. We refer to these additional days as “staircase”. After 1 -5 days of staircase training, we stopped providing cutaneous stimulation and began the washout portion of the experiment. We used this experimental approach for both prelesion and postlesion experiments in our LMAN, 6-OHDA, and Sham data sets. Importantly, although the number of days of staircase varied between individual birds, for each individual bird we matched the same number of prelesion days of staircase and postlesion days of staircase to ensure that, in both experimental conditions, the bird had an equivalent amount of time and opportunity to learn.

Custom-written MATLAB software (The MathWorks) was used for song analysis. On each day of every experiment, we quantified important song features, such as the pitch, amplitude, and spectral entropy, of all renditions of the targeted syllable produced between 10 A.M. and 12 P.M. We did so to account for potential circadian effects on song production. To ensure a level of consistency in number of target syllable renditions measured on each day of an experiment, and to have a minimum number of syllable renditions necessary to get an accurate measure of average syllable pitch, we checked that at least 30 renditions of the target syllable were sung within the 10 A.M. to 12 P.M. window. If there were less than 30 renditions of the target syllable, we extended the time window for song analysis by 1 hour in both directions (9 A.M. to 1 P.M.) and then reassessed to see if there were at least 30 syllable renditions. If not, we continued this process of extending the time window by 1 hour until 30 song renditions were in that day’s data set. Daily targeting sensitivity (hit rate) and precision (1-false-positive rate) were measured in all experiments to ensure accurate targeting of the specific target syllable (and not accidentally targeting different song syllables). During the pitch-contingent feedback portion of the experiment, a subset (10%) of randomly selected target syllables did not trigger feedback, regardless of syllable pitch. These “catch trials” allowed for the quantification and comparison of syllable features, such as pitch, amplitude, and entropy between trials when feedback was provided and trials when feedback was not provided. Pitch changes were quantified in units of semitones as follows:

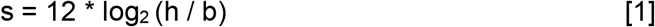

where s is the pitch change (in semitones) of the syllable, h is the average pitch (in Hertz) of the syllable, and b is the average baseline pitch (in Hertz) of the syllable.

### Analysis of Variability in Syllable Pitch

We compared pitch variability pre-and postlesion using methods described in prior literature^30–32^. We analyzed all song renditions (within the 10 A.M. -12 P.M. time window) performed on the final day of baseline prelesion and on the final day of baseline postlesion. We did so in our LMAN lesion experimental group as well as our 6-OHDA lesion experimental group. To measure the variability in pitch of the song syllables, we calculated the coefficient of variation (CV) for the pitch of each syllable using the following formula: CV = (Standard Deviation / Mean) * 100.

### LMAN Lesions

Birds were anesthetized under ketamine and midazolam and were mounted in a stereotax. The beak angle was set to 20° relative to the surface level of the surgery table. For stereotactic targeting of specific brain regions (in this case, LMAN), anterior-posterior (AP) and medial-lateral (ML) coordinates were found relative to Y_0_, a visible anatomical landmark located at the posterior boundary of the central venous sinus in songbirds. Dorsal-ventral (DV) coordinates were measured relative to the surface of the brain. Bilateral craniotomies were made at the approximate AP coordinates 4.9 mm to 5.7mm and ML coordinates 1.5 mm to 2.5 mm. A lesioning electrode was then inserted 1.9 mm to 2.1 mm below the brain surface. These stereotactic coordinates targeted locations within LMAN. We then passed 100 μA of current for 60-90 seconds at 5-6 locations in LMAN in both hemispheres in order to electrolytically lesion the areas. This methodology was based on prior work involving LMAN lesions and LMAN inactivations^20,26,30–32,49^. In sham operated birds, we instead performed small lesions in brain areas dorsal to LMAN. Again, this was consistent with methodology from prior studies^20,30,31^.

Birds recovered within two hours of surgery and began singing normally (at least 30 renditions of target syllable within 2 hours) typically 3 to 8 days after surgery.

Behavioral measures indicated that LMAN was effectively lesioned in the birds in the LMAN lesion data set. LMAN lesions in adult songbirds produce a significant decrease in the trial-to-trial variability of song syllable pitch^30–32^. To assess lesion-induced changes in the variability of syllable pitch between conditions (LMAN lesion and sham), we calculated the CV of syllable pitch pre-and postlesion. We found that LMAN lesions induced a significant decrease in pitch CV (Fig. 3b; paired t-test,). Sham operations did not induce a significant change in syllable CV (Fig. 3b; paired t-test, p = 0.911). The lesion-induced changes in syllable CV (post -pre) were significantly greater than changes to CV in sham lesioned controls (Figure 3-Figure Supplement 1a; 2 sample KS test, p = 0.003).]

Lesions were confirmed histologically using cresyl violet staining after completion of behavioral experimentation. In tissue from sham operated birds, we identified Area X and LMAN based on regions of denser staining as well as well-characterized anatomical landmarks^50^. The histology methodology we employed followed previous literature involving LMAN lesions^20,30^. We performed Nissl stains to stain for neuronal cell bodies in brain slices after experiments were complete (Figure 3-Figure Supplement 2a). We then calculated the optical density ratio of the region containing LMAN compared to background (a pallial region outside of LMAN) (Figure 3-Figure Supplement 2b)^22,29^. The distribution of OD ratios from LMAN lesioned tissue was significantly less than the OD ratios from sham lesioned tissue (Figure 3-Figure Supplement 2c; 2 sample KS test, p < 0.0010). This suggests that the density of neuronal cell bodies within LMAN was reduced following electrolytic lesions compared to following sham. Similar to a prior study, we also qualitatively assessed each slice of brain tissue to measure the percentage of intact LMAN remaining in the tissue^20^. We found that all of the LMAN lesioned birds had 80-100% of LMAN lesioned in both hemispheres.

### 6-OHDA Lesions

Birds were anesthetized using ketamine and midazolam and were mounted in a stereotax, where the beak angle was set to 20° relative to the surface level of the surgery table. Isoflurane was used in later hours of the surgery to maintain an anesthetized state. Bilateral craniotomies were made above Area X from the approximate AP coordinates 4.5 mm to 6.5mm and ML coordinates 0.75 mm to 2.3 mm relative to Y_0_.

In each hemisphere, we inserted a glass pipette containing a 6-OHDA solution (see below) and made 12 pressure-injections in a 3 mm x 4 mm grid between AP coordinates 5.1 mm and 6.3 mm, ML coordinates 0.9 mm and 2.2 mm and the DV coordinate 3.18 mm relative to Y_0_. Additional bilateral 6-OHDA injections were made at the AP coordinate 4.8 mm, ML coordinate +/-0.8 mm, and DV coordinate 2.6 mm from the brain surface to lesion the most medial portion of Area X. Each injection consisted of 13.8 nL of 6-OHDA solution, injected at a rate of 23 nL/s at each site. The pipette was kept in place for 30 seconds after each injection and was then slowly removed. In sham operated birds, we performed the same surgical operations, except saline was injected into Area X instead of 6-OHDA. Again, this was consistent with methodology from prior studies^22,29^.

Birds recovered within two hours of surgery and began singing normally (at least 30 renditions of target syllable within 2 hours) typically 3 to 8 days after surgery. 6-OHDA solution was prepared using 11.76 mg 6-OHDA-HBr and 2 mg ascorbic acid in 1 mL of 0.9% normal saline solution. The solution was light-protected after preparation to prevent oxidation.

In order to confirm the effectiveness of 6-OHDA injections at lesioning dopaminergic input to Area X, we quantified the extent of the reduction of catecholaminergic fiber innervation within Area X after completing the behavioral experimentation in each bird^22,29^. To visualize dopaminergic innervation, we labeled tissue with a common biomarker for catecholaminergic cells (Figure 4-Figure Supplement 1a). To determine whether the concentration of dopaminergic fibers in Area X had decreased, we measured the optical density ratio (OD): the ratio of the stain density of Area X to the stain density of the surrounding striatum. OD ratios from individual 6-OHDA lesioned brains decreased compared to control (Figure 4-Figure Supplement 1b). The distribution of all OD ratios from all of the 6-OHDA lesioned tissue was significantly lower than that of the brain tissue from sham operated birds (Figure 4-Figure Supplement 1c; 2 sample KS test, p < 0.001). These results are similar to previous reports that used 6-OHDA injections to lesion dopaminergic input to Area X^22,29^, and they indicate that the 6-OHDA injections successfully lesioned dopaminergic input to Area X.

Lesion size was quantified by determining the proportion of 6-OHDA lesioned tissue that had an OD ratio of Area X to non-X striatum that was less than the fifth percentile of OD ratios in sham tissue. There was not a significant correlation between lesion size and the lesion-induced change in learning magnitude (post-pre) (Figure 4-Figure Supplement 2a, b; R^2^ = 0.019, p = 0.137).

### Histology

Between 14 and 54 days after surgery, birds were injected with a lethal dose of ketamine and midazolam and were perfused. Tissue was post-fixed in 4% paraformaldehyde at room temperature for 4-16 hours and then moved to a solution of 30% sucrose for at least one day at 4°C for cryoprotection. Then, brain tissue was sliced in 40 μm sections. A chromogenic tyrosine hydroxylase (TH) stain was used to quantify the depletion of catecholaminergic fiber innervations in tissue collected from 6-OHDA lesioned birds, and Nissl and fluorescent NeuN staining was used to assess the density of cell bodies in tissue from LMAN lesioned and sham operated birds. For one bird in the 6-OHDA lesioned group, a Nissl stain was performed on alternate tissue sections to ensure no cell death occurred as a result of the lesion.

For TH immunohistochemistry, tissue was incubated overnight in a primary anti-TH antibody solution. The tissue was next incubated in biotinylated horse anti-mouse secondary antibody solution for 1 hour. Then, the tissue was submerged in a diaminobenzidine (DAB) solution (2 DAB tablets, Amresco E733 containing 5 mg DAB per tablet, 20 mL Barnstead H_2_O, 3 μL H_2_O_2_) for less than 5 minutes for visualization. The DAB solution was prepared 1h prior to use. Tissue was washed, mounted and coverslipped using Permount mounting medium.

### Tyrosine Hydroxylase Stain

Between each incubation tissue was washed with 0.1 M phosphate buffer (PBS) (23 g dibasic sodium phosphate, 5.25 g monobasic sodium phosphate, and 1 L deionized H_2_O) 3 times for 10 min each. Tissue was first washed and then incubated in 0.3% H_2_O_2_ for 30 min and then 1% NaBH_4_ for 20 min, followed by overnight incubation in a primary anti-tyrosine hydroxylase antibody solution. The tissue was next incubated in biotinylated horse anti-mouse secondary antibody solution for 1 h, then incubated in avidin-biotin-complex (ABC) solution for 1 h that had been prepared 30 min prior to use. The tissue was then submerged in a diaminobenzidine (DAB) solution for less than 5 minutes. Tissue was then washed, mounted and coverslipped using Permount mounting medium. These TH stains mark neurons expressing TH, which are catecholaminergic.

### Nissl Stain

Tissue was washed in 0.1 M PBS three times for 10 minutes and was then mounted. The slides were incubated in Citrisolv twice for 5 min each, then delipidized in the following ethanol concentrations for two minutes each: 100%, 100%, 95%, 95%, and 70%. The tissue was briefly (less than 15 s) rinsed in deionized water, then was incubated in cresyl violet (665 μL glacial acetic acid, 1 g cresyl violet acetate, and 200 mL deionized water) for 30 min. The tissue was rinsed in deionized water, then briefly (less than 15 seconds) submerged in the following ethanol concentrations for 2 min each: 70%, 95%, 95%, 100%, and 100%. The tissue was then incubated in citrisolv twice for 5 min. The tissue was coverslipped using Permount mounting medium. These Nissl stains mark neuronal cell bodies.

### NeuN Antibody Stain

Between each incubation, tissue was washed with 0.1 M PBS 3 times for 10 min each. Tissue was incubated in primary antibody solution (4 mL EMD Millipore guinea pig anti-NeuN Alexa Fluor 488 antibody, 6 mL Triton X-100, 20 mL normal donkey serum (NDS) and 1.95 mL 0.1 M PBS) overnight. The tissue was then washed and incubated in a secondary antibody solution (10 mL Jackson Labs donkey anti-guinea pig (DAG), 6 mL Triton X-100, and 1.975 mL 0.1 PBS) overnight. Tissue was then washed, mounted and coverslipped with Flurogel mounting medium. Slides were sealed with lacquer. Images were taken under a widefield microscope (BioTek Lionheart FX, Sony ICX285 CCD camera, Gen5 acquisition software, 1.25x magnification, 16-bit grayscale).

### Lesion Analysis

Analysis of lesions was based on previously published methodology^22,29^. Images of stained tissue sections were obtained using a slide scanner and were converted into 8-bit grayscale images in ImageJ. In control (unoperated) birds, Area X stains darker than surrounding striatum in TH-DAB-stained tissue due to a higher density of catecholaminergic inputs in Area X^22,29^. The baseline level of stain darkness can vary from bird to bird. Therefore, rather than directly comparing the stain density of lesioned and sham tissue, the ratio of the stain density of Area X to that of the surrounding striatum (OD ratio) was calculated to determine whether the concentration of catecholaminergic fibers was decreased. Prior work demonstrated that the vast majority of catecholaminergic input to Area X is dopaminergic^22^.

For each section of tissue containing Area X, a customized ImageJ macro was used to select regions of interest (ROIs) within Area X and within a portion of striatum outside Area X by manually outlining Area X and selecting a circular 0.5 mm-diameter region of striatum anterior to Area X. Pixel count and optical density (OD) of each ROI were measured, and the density of TH-positive fibers was calculated using the ratio of the OD of Area X to the OD of non-X-striatum.

The cumulative distribution of OD ratios for sham operated birds was used to construct a 95% confidence interval and determine the threshold for lesioned tissue. 6-OHDA-lesioned tissue in which the OD ratio fell below the 5th percentile of control tissue had a significantly reduced TH-positive fiber density.

### Statistical Testing

All error bars presented in the main text represent SEM. When assessing whether a significant amount of vocal learning occurred in one experiment, we used one-sample t-tests to compare the mean pitch on the final day of training vs zero. To assess whether a significant difference in amount of learning occurred within an individual bird pre-vs postlesion, we used paired t-tests. To assess significance between distributions of target syllable pitches on various days of the experiment (Baseline, shock, washout), we used a 2-sample KS test.

Each experimental group had at least five birds, and for each bird, the target syllable was typically repeated well over 30 times a day. Therefore, the structure of our data is hierarchical, so error accumulates at different levels (birds and syllable iterations). Simply grouping all the data together ignores the non-independence between samples and underestimates the error. To address this issue, we employed a hierarchical bootstrap method to measure SEM and calculate p-values^28^. For each experimental day we calculated normalized pitch values (in semitones) (normalized to the mean pitch on the final baseline day during that particular experiment). We then generated a population of 10000 bootstrapped means according to the following sampling procedure: to generate each individual subsample, we resampled across each level of hierarchy in our data (first resampled among the birds, then for each selected bird, we resampled among syllable iterations). The standard deviation of this population of bootstrapped means provides an accurate estimate of the uncertainty of the original data^28,29^. Thus, the SEM values (which are used for error bars) we report when employing the hierarchical bootstrap method are equal to this standard deviation.

To calculate p-values and determine significance for comparing our data to zero using the hierarchical bootstrap method, we calculated P_boot_: the proportion of bootstrapped means greater than zero compared to the total number of bootstrapped means. Using an acceptable type-1 error rate of .05, any value of this P_boot_ ratio greater than .975 indicates the mean was significantly greater than zero and any value less than .025 indicates the mean was significantly less than zero. P_boot_ values between .025 and .975 indicate no significant difference between the data set and zero. Because we measure adaptive pitch changes in semitones, which are a normalized measure of pitch change where baseline is set to zero, this method of calculating P_boot_ was employed in all instances where it was necessary to assess whether there was a significant change in pitch at the end of training compared to baseline (zero).

We also sometimes sought to determine significance for the comparison of two means rather than what was previously described (where we assess significance between one mean compared to baseline (zero). We used a similar hierarchical bootstrap statistical methodology and calculated P_boot_. The key difference is that, rather than measuring the proportion of resampled means greater than or less than zero, we instead calculate a joint probability distribution for the means of the two resampled data sets. We measured the percentage of this joint probability distribution that was above one side of the unity line. This percentage is the P_boot_ values we report in these instances. If the proportion of this joint probability distribution that falls above the unity line is greater than .975, it indicated a significantly greater mean of data set 1 over data set 2. If the percentage of the joint probability distribution that was above the unity line was less than .025, it indicated a significantly lower mean of data set 1 compared to data set 2. P_boot_ values between .025 and .975 indicate no significant difference between the two data sets. This method was employed in all instances where it was necessary to assess whether the learning magnitudes (adaptive pitch changes by the end of training) were significantly different pre-vs postlesion (or pre-vs postsham) or across experimental conditions (e.g., postsham vs postlesion or post LMAN lesion vs post 6-OHDA lesion).

In both forms of P_boot_ calculation, the lowest statistical limit for P_boot_ is P_boot_ <0.0010, due to resampling 10^4^ times to create bootstrapped means. The highest possible limit for P_boot_ is P_boot_ > 0.9999, for the same reason.

Sample sizes were not predetermined using a power analysis. Sample sizes of all sets of experiments were comparable to relevant prior literature^22,27,29^. If at any point during cutaneous stimulation training or white noise training a bird’s rate of singing dropped below 10 songs per day for over one day, that experiment was stopped and the data were excluded from further analysis.

## Supplementary Figures

**Figure 2-Figure Supplement 1.**
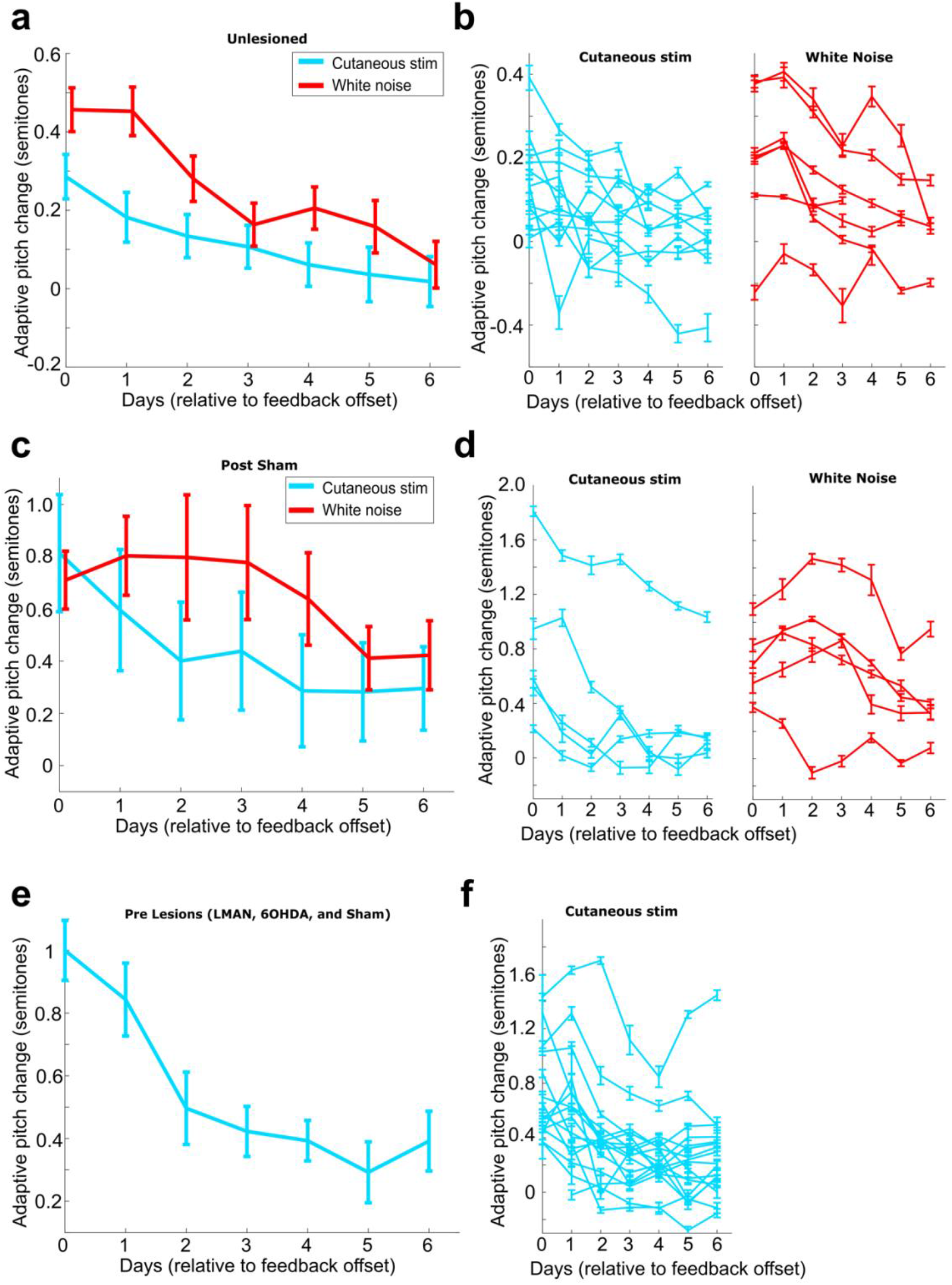
Rates of washout across different experimental conditions. (**a**) Adaptive pitch change (measured relative to baseline) during washout from the group of birds who received no invasive brain operations (n=13 experiments). Adaptive pitch change did not significantly differ between white noise and cutaneous stimulation training experiments on any of the days of washout (0.487 < P_boot_ < 0.541 on each day of washout, where 0.025 < P_boot_ < 0.975 indicates no significant difference between means). (**b**) The same washout data from (a), except each trace is the data from an individual experiment. (**c**) Adaptive pitch change (measured relative to baseline) during washout from the sham lesioned data set (n=6 experiments). Adaptive pitch change did not significantly differ between white noise and cutaneous stimulation training experiments on any of the days of washout (0.370 < P_boot_ < 0.900) (**d**) The same washout data from (c), except each trace is the data from an individual experiment. (**e**) Adaptive pitch change (measured relative to baseline) during washout from all prelesion experiments in birds who received invasive brain operations (presham, pre LMAN lesion, and pre 6-OHDA lesion), n = 16. (**f**) The same washout data from (e), except each trace is the data from an individual experiment.

**Figure 2-Figure Supplement 2.**
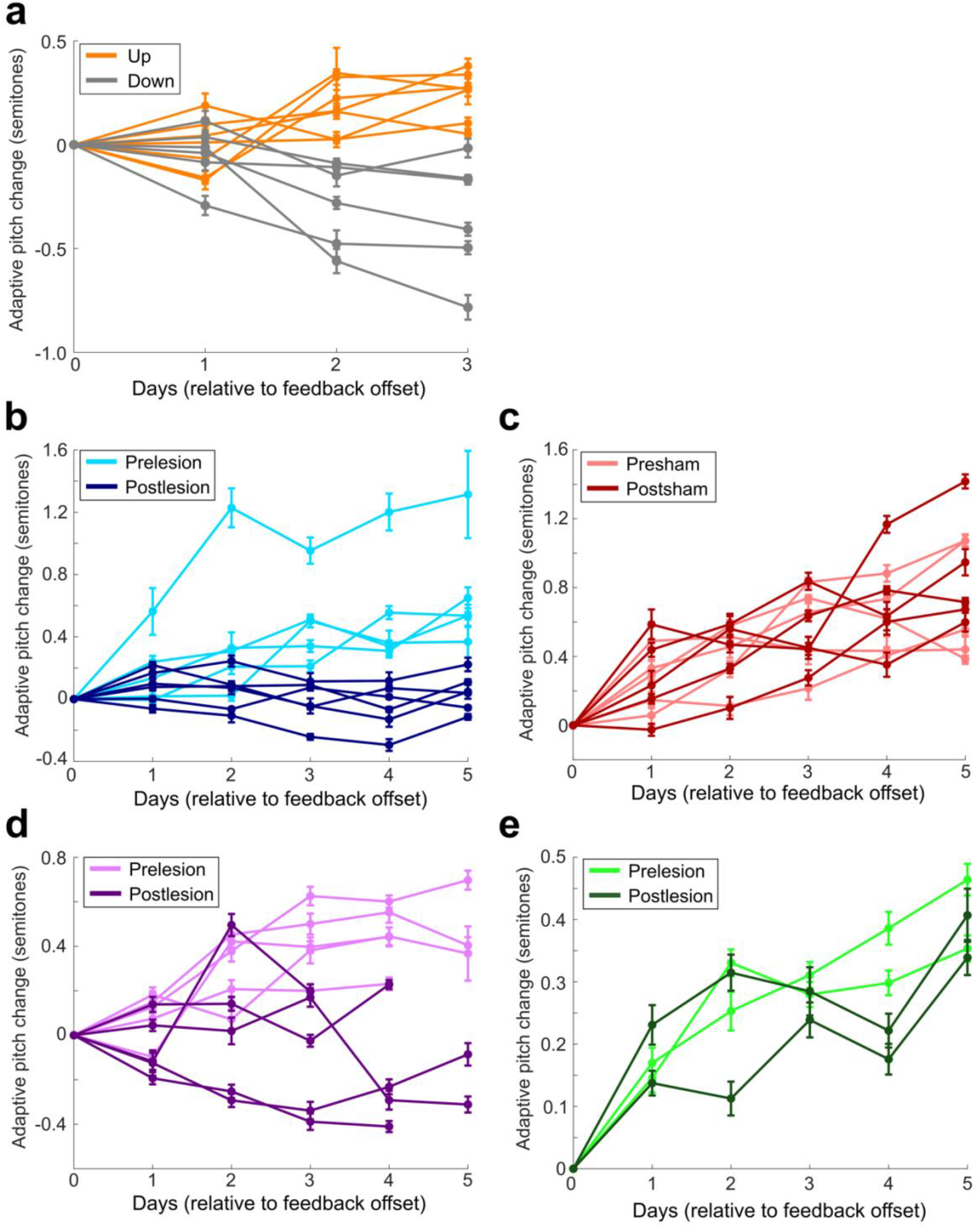
Amount of pitch change on each day of cutaneous stimulation training for each individual experiment. (**a**) All experiments performed in birds who did not undergo any invasive brain operations (LMAN lesions, 6-OHDA dopamine lesions, sham operations). Orange are experiments where upwards pitch change resulted in less frequent triggering of cutaneous stimulations. Gray are experiments where downwards pitch change resulted in less frequent triggering of cutaneous stimulations. (**b**) Results from all experiments performed in birds who underwent LMAN lesions. (**c**) Results from all experiments performed in birds who underwent LMAN sham operations. (**d**) Results from all experiments performed in birds who underwent 6-OHDA lesions. (**e**) Results from all experiments performed in birds who underwent 6-OHDA sham operations.

**Figure 2-Figure Supplement 3.**
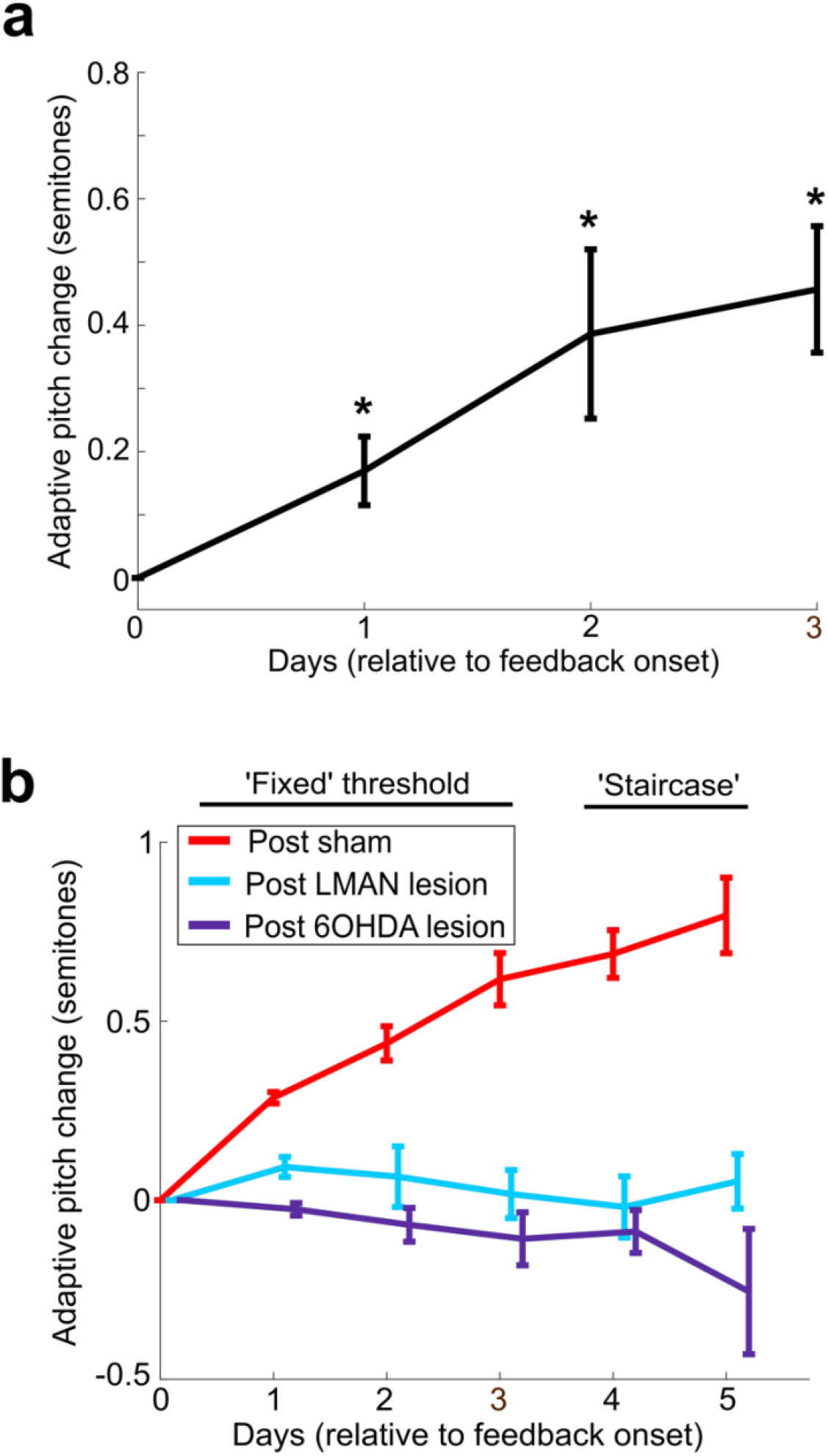
(**a**) Adaptive change in target syllable pitch (in semitones) during three days of white noise training in 8 birds who did not undergo any lesions or sham operations. Probability of resampled mean pitch on each day of training lesser than or equal to zero was Pboot<0.0010. (**b**) Learning magnitudes (adaptive change in target syllable pitch in semitones) during five days of white noise training in birds who underwent sham operations, LMAN lesions, and 6-OHDA lesions. Only postsham learning magnitude was significantly greater than baseline (probability of resampled mean pitch on each of the final four days of training lesser than or equal to zero was Pboot<0.0010). Post LMAN lesion learning magnitudes were significantly less than postsham (probability of resampled mean pitch of post LMAN lesion data on the final four days of training lesser than or equal to resampled mean pitch of postsham data was Pboot<0.0010).

**Figure 2-Figure Supplement 4.**
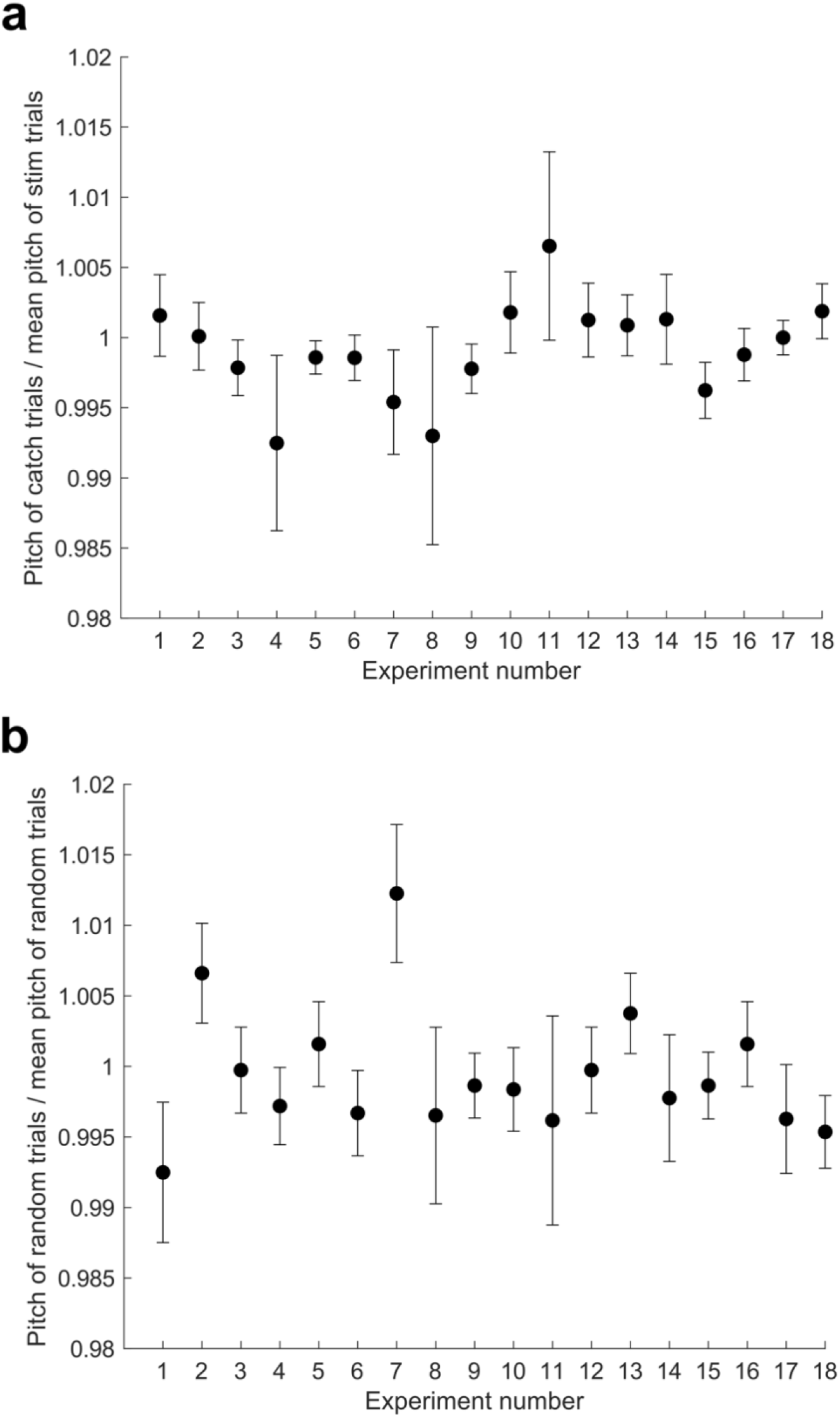
Analysis of acute effects of cutaneous stimulation on target syllable pitch (**a**) For each experiment throughout all data sets described in this paper, we calculated the pitch of every catch trial that occurred during each day of cutaneous stimulation training, normalized to the mean pitch of all trials that triggered cutaneous stimulations. We excluded all experiments with less than 10 catch trials. Error bars are S.E.M. No individual experiment differed significantly from 1 (t-test, 0.071 < p < 0.997). (**b**) Same as in (a), but we analyzed randomly selected trials from a baseline recording day for each experiment. For each experiment, we selected the same number of trials as in (a). There was no significant difference between this data set and the normalized catch trials (paired t-test, p = 0.339).

**Figure 3-Figure Supplement 1.**
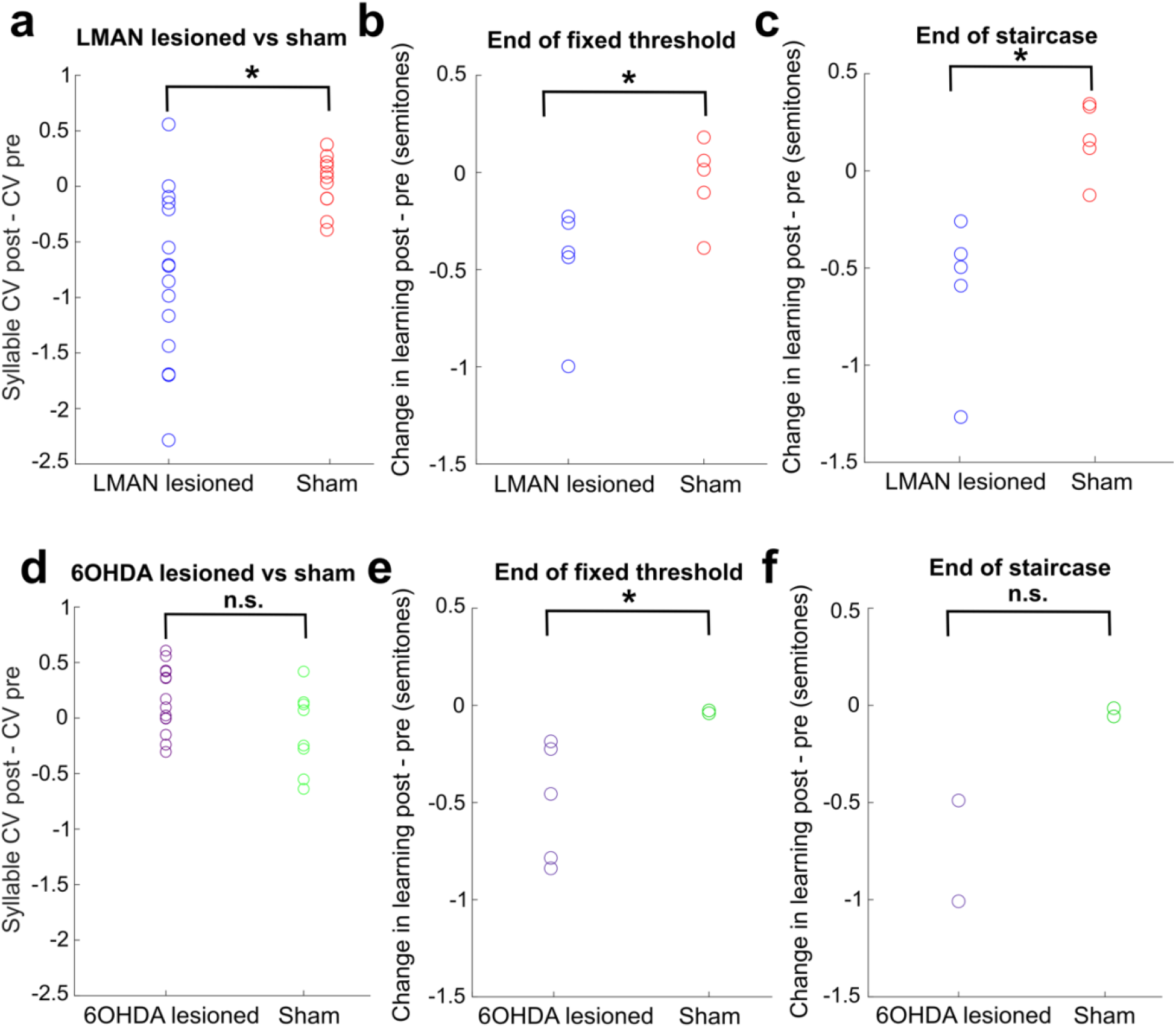
(**a**) Change in syllable CV in LMAN lesioned and sham operated birds. Each data point represents the CV postlesion - CV prelesion of one individual song syllable. LMAN lesions induced a significant reduction in syllable CV compared to sham operations (2 sample KS test, p=0.003). (**b**) Lesion-induced change in learning magnitude (measured at the end of three days of cutaneous stimulation training) in LMAN lesioned and sham operated birds. The lesion-induced change in learning magnitude (post–pre) for LMAN lesioned birds was significantly greater than sham (2 sample KS test: p=0.036). (**c**) Same as (b), except learning magnitude was measured at the end the extended staircase portion of training. The lesion-induced change in learning magnitude (post – pre) in LMAN lesioned birds was significantly greater than in sham operated birds (2 sample KS test, p=0.004) (**d**) Change in syllable CV in 6OHDA lesioned and sham operated birds. Each data point represents the CV postlesion -CV prelesion of one individual song syllable. 6OHDA lesions did not induce a significant reduction in syllable CV compared to sham operations (2 sample KS test, p=0.209). (**e**) Lesion-induced change in learning magnitude (measured at the end of three days of training) in 6OHDA lesioned and sham operated birds. The lesion-induced change in learning magnitude (post–pre) in 6OHDA lesioned birds was significantly greater than in sham operated birds (2 sample KS test: p=0.036). (**f**) Same as (e), except learning magnitude was measured at the end the staircase portion of training.

**Figure 3- Figure Supplement 2.**
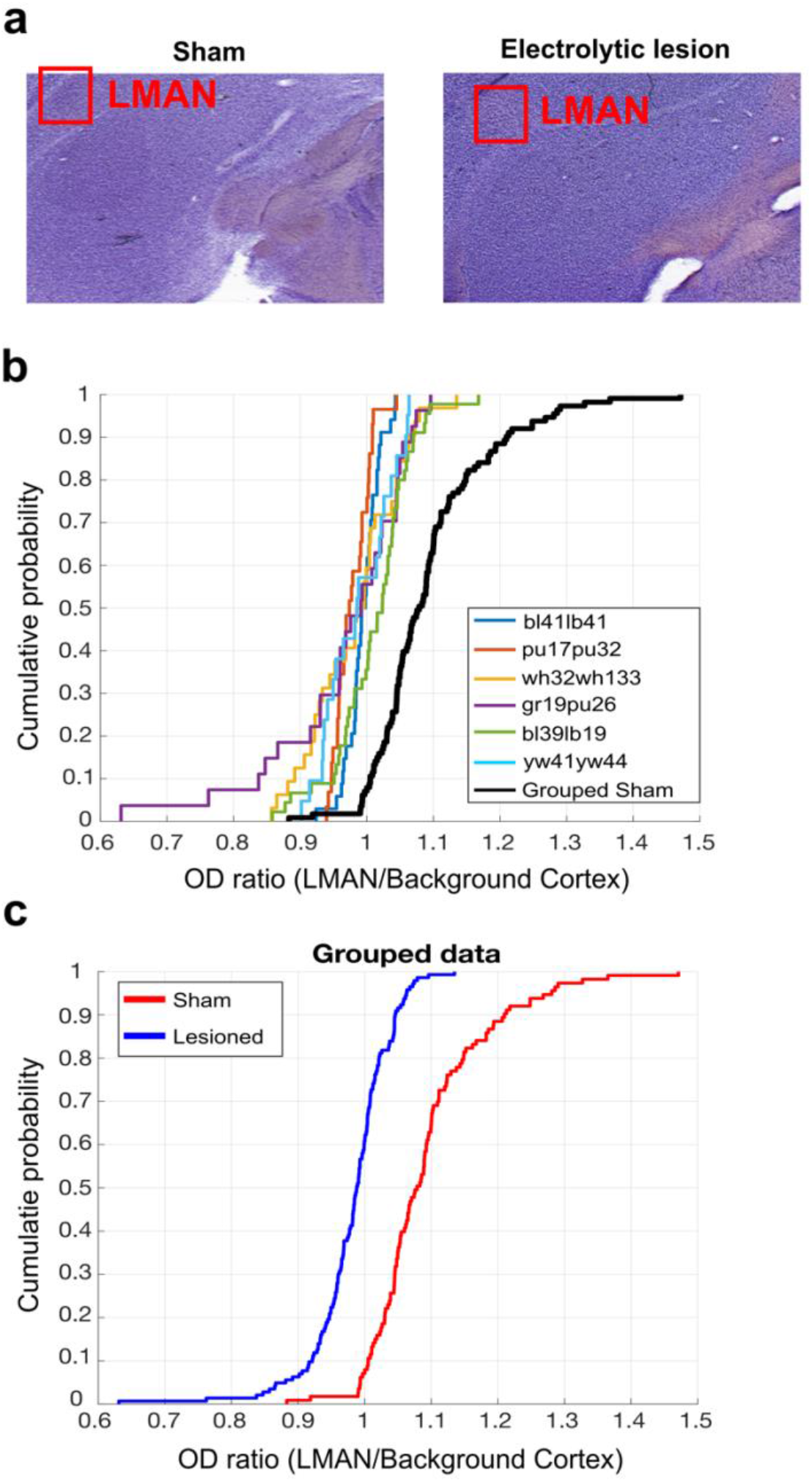
LMAN lesion histological analysis. (**a**) Example images of Nissl-stained brain tissue. Tissue from sham operated bird on the left and tissue from LMAN lesioned bird on the right. Red boxes highlight the locations of LMAN. (**b**) CDF plot of optical density (OD) ratios (OD of LMAN / OD of non-LMAN-pallium) in lesioned and sham operated birds. Each line shows the OD ratios from each individual LMAN lesioned bird, and the black line shows the OD ratios from the grouped sham data set. (**c**) CDF plot of OD ratios in lesioned and sham operated birds. Blue line shows the OD ratios from the grouped LMAN lesion data set and the red line shows the OD ratios from the grouped sham data set (2 sample KS test, p < 0.001).

**Figure 4- Figure Supplement 1.**
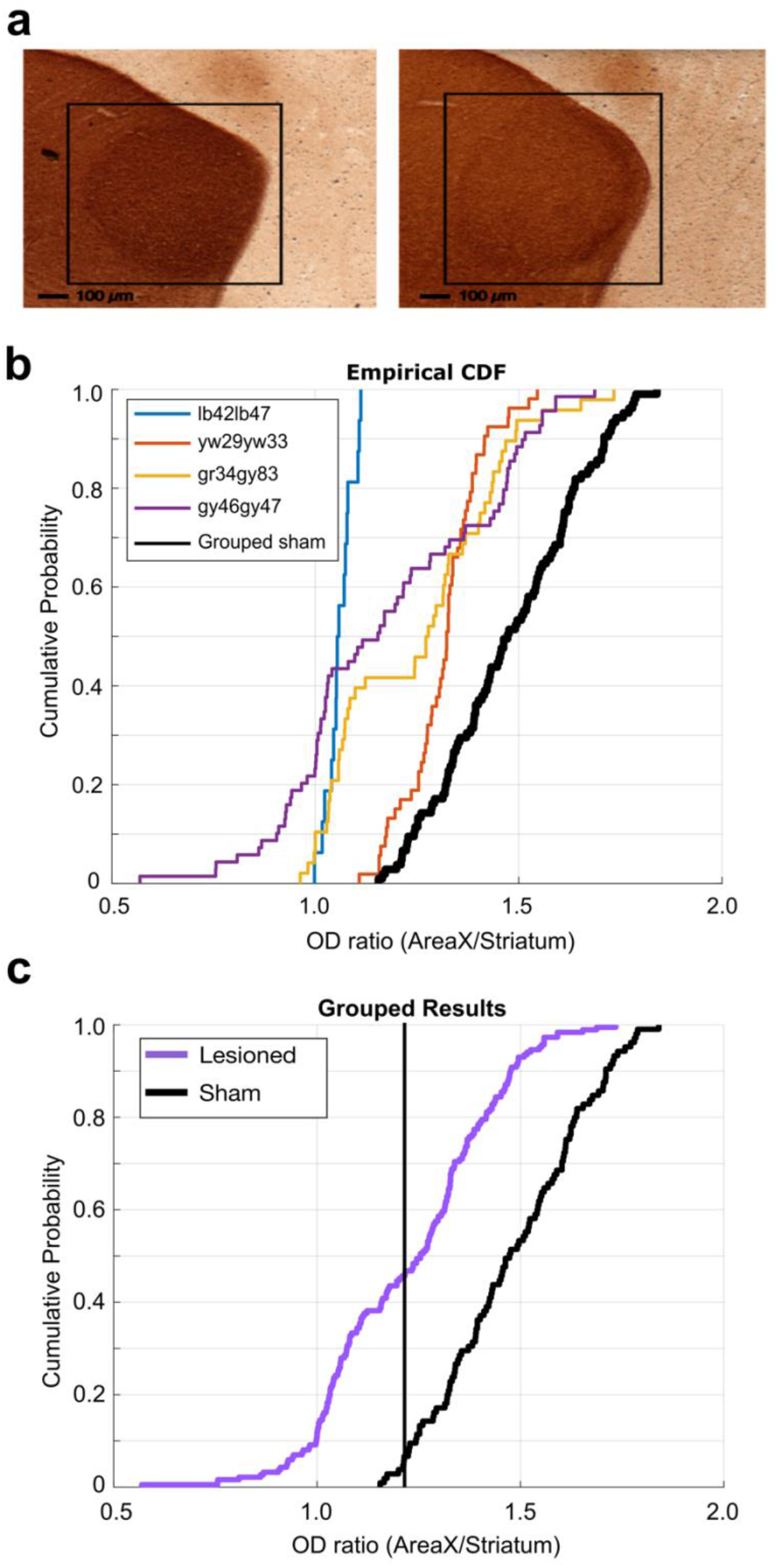
6-OHDA lesion histological analysis. (**a**) Example images of TH-stained brain tissue. Tissue from sham operated bird on the left and tissue from 6-OHDA lesioned bird on the right. Black boxes highlight the locations of Area X. (**b**) Cumulative probability plot of optical density (OD) ratios (OD of Area X / OD of non-X-striatum) in 6-OHDA lesioned and sham operated birds. Each line shows the OD ratios from each individual 6-OHDA lesioned bird, and the black line shows the OD ratios from the grouped sham data set. (**c**) CDF plot of OD ratios in lesioned and control birds. Purple line shows the OD ratios from the grouped 6-OHDA lesioned dataset, and the black line shows the OD ratios from the grouped sham dataset (2 sample KS test, p < 0.001).

**Figure 4- Figure Supplement 2.**
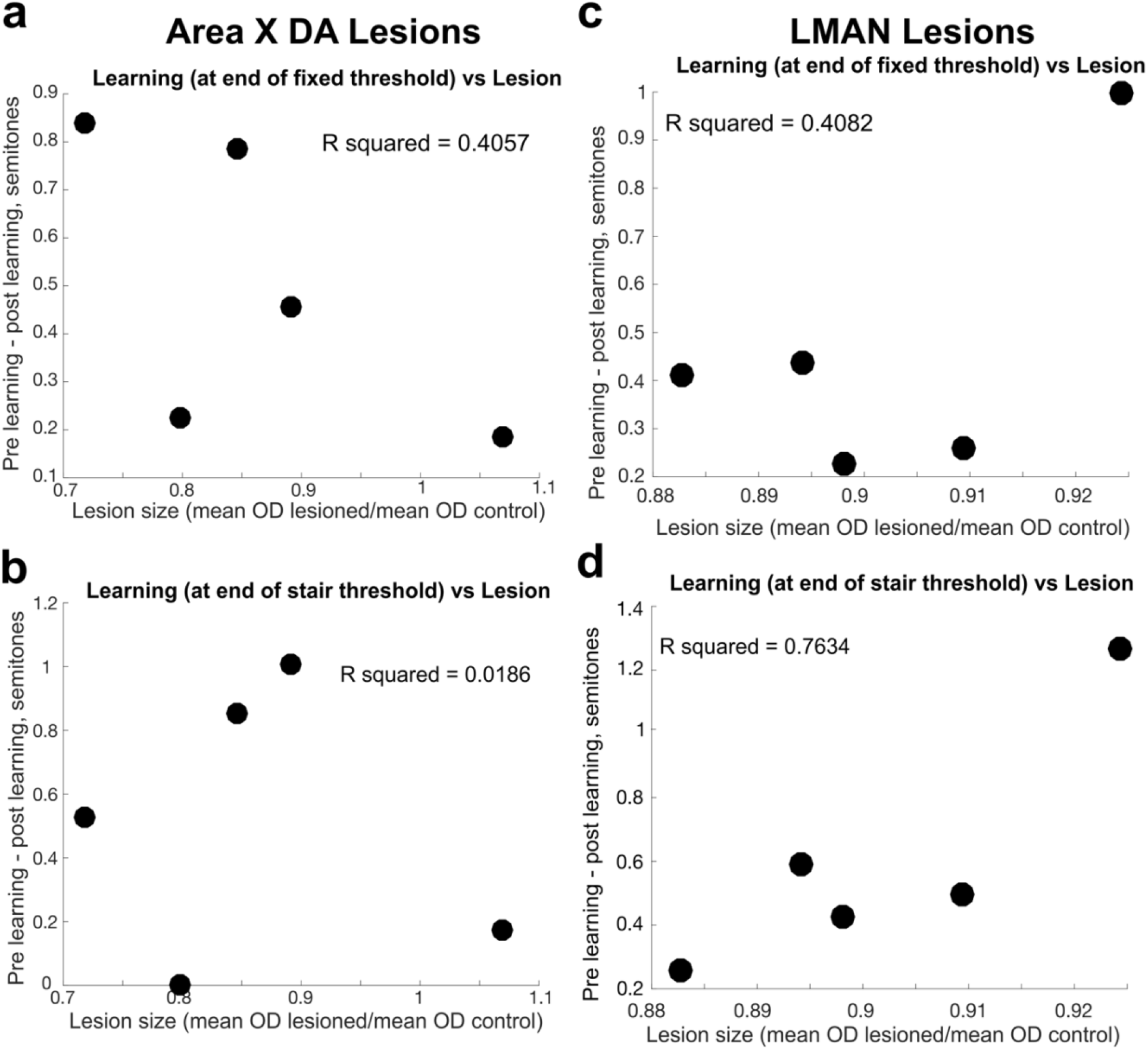
Comparison of lesion magnitude and learning deficit. (**a**) For each bird, the difference between the magnitude of learning, calculated at the end of three days of cutaneous stimulation training, prelesion vs postlesion, compared to the magnitude of the Area X dopamine lesion in 6-OHDA injected birds, measured by the ratio of the mean OD of the lesioned tissue to the mean OD of control tissue. Each dot represents the results from each individual bird. (**b**) Same as in (a), but the magnitude of learning was assessed at the end of the additional days of staircase training. (**c**) For each bird, the difference between the magnitude of learning prelesion and the magnitude of learning postlesion (in both cases, the magnitude of learning is measured at the end of the three days of fixed threshold training, compared to the size of the LMAN lesion in electrolytically lesioned birds, measured by the ratio of the mean OD of the lesioned tissue to the mean OD of control tissue. Each dot represents the results from each individual bird. (**d**) Same as in (c), but the magnitude of learning was assessed at the end of the additional days of staircase training.

